# Macrogenetic atlas of prokaryotes reveals selection-driven structures

**DOI:** 10.1101/2025.02.26.640246

**Authors:** Chao Yang, Fuxing Jiao, Naike Wang, Shanwei Tong, Hao Huang, Wenxuan Xu, Xavier Didelot, Ruifu Yang, Yujun Cui, Daniel Falush

## Abstract

Macrogenetics investigates patterns and predictors of intraspecific genetic variation across diverse taxa, offering a framework for understanding species diversity and addressing evolutionary hypotheses. Here, we present a macrogenetic atlas of prokaryotes (MAP), integrating genomic data from 15,235 prokaryotic species (30 parameters in 12 categories) and population genetic data from 786 species (22 parameters in 7 categories), together with phylogenetic, phenotypic, and ecological information. MAP enables quantitative species characterization, straightforward cross-species comparisons, and easy generation, testing, and refinement of evolutionary hypotheses. For example, our analyses show that long- and short-range genetic linkage capture distinct evolutionary dynamics and independently shape genetic diversity. Further, we propose that as within-species diversity increases, epistatic selection increasingly contributes to shaping genetic diversity, exemplified by convergent selection-driven structures (ecospecies) identified in *Streptococcus mitis* and *S. oralis*. Overall, MAP represents a widely applicable resource (www.genomap.cn) and offers novel insights into the drivers of macroevolution and the life cycle of prokaryotic species.

## Introduction

Life is diverse, and its understanding requires macro-scale frameworks that capture species diversity, akin to maps guiding exploration. Since Linnaeus, taxonomy has provided this foundation based on phenotypic classification, and is now being systematically updated by the addition of genomic data, for example in the Tree of Life project^1^. The integration of genotype, phenotype, and ecological data remains the major unresolved challenge and is essential for characterizing species and identifying the forces driving diversification and speciation. Within microbiology, classical resources such as *Bergey’s Manual*, which integrates taxonomy with phenotypic and ecological information^2^, have long served this purpose.

Incorporating within-species genomic variation data is the next critical step. Advances in high-throughput sequencing have produced extensive genomic resources across many taxa^3,4^. With their rapid accumulation, the time is now ripe for systematic integration under a new framework, macrogenetics, which embraces broad taxonomic, spatial and temporal scales across dozens to thousands of species^5,6^. Macrogenetics provides a framework for exploring species diversity and testing long-standing evolutionary hypotheses. For example, community-level species richness has been shown to correlate positively with within-population genetic diversity in fish and mammals^7,8^, supporting the species-genetic diversity correlation (SGDC) concept^9^. Other studies have linked genetic diversity to life-history traits such as fecundity and propagule size^10^, and revealed latitudinal gradients in genetic diversity^11^.

Prokaryotes make up more biomass than all other organisms except plants^12^. Furthermore, a single species often contains lineages adapted to multiple niches, differing in host range, pathogenicity and nutrient use, as exemplified by *Escherichia coli*^13^. This contrasts with eukaryotes, where within-species differentiation and specialization are typically limited. Such pervasive intraspecific diversity in prokaryotes raises fundamental questions: how are ecological niches partitioned, and how do phenotypic and niche diversity shape genomic and population genetic features? These differences also imply that differentiation and speciation in prokaryotes are driven by forces distinct from those shaping eukaryotes^14^, but these remain poorly understood.

Here we present the Macrogenetic Atlas of Prokaryotes (MAP), an open resource (www.genomap.cn) that integrates large-scale genomic and population genetic data with phylogenetic, phenotypic and ecological information. MAP assigns each species a quantitative “position” based on ranked features across species, enabling quantitative characterization and straightforward cross-species comparisons. It also serves as an empirical resource for generating, testing, and refining evolutionary hypotheses. Overall, MAP provides a framework for exploring and understanding prokaryotic diversity.

## Results

### Overview of MAP

We analyzed 317,542 assembled genomes across 15,235 species (503 archaeal and 14,732 bacterial; NCBI taxonomy). After deduplication and quality control, we generated representative (*Rep.*) datasets for each species to reduce repeat-sampling biases, and clonal group (CG) datasets to capture microevolution (**Fig. 1**; Methods; Supplementary Tables 1-2, Supplementary Fig. 1).

**Figure 1.**
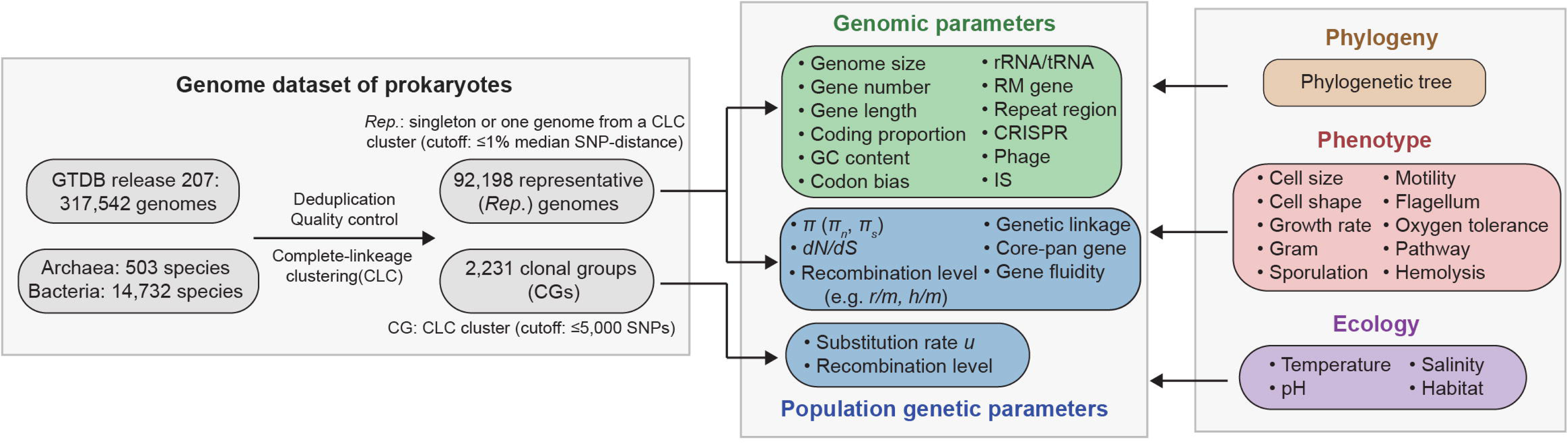
**Study flowchart of the MAP.**

Species-level genomic parameters (GPs) were defined as the median across *Rep.* genomes, whereas population genetic parameters (PGPs) were estimated for species represented by ≥10 genomes, using random samples of ten *Rep.* genomes, CG datasets, or all *Rep.* genomes (Methods). To ensure robustness, only PGPs with low variability (standard deviation/median < 0.5) across random-sampling replicates were retained, yielding estimates highly consistent with full datasets (correlation coefficient *r* > 0.8 for all but one parameter; *h/m*, *r* = 0.68) and independent of sample size (Supplementary Figs. 2-4). By contrast, all GP and CG-based PGP estimates were retained across species to maximize cross-species comparability, with high variability reflecting substantial within-species heterogeneity.

In total, we obtained 30 GPs across 12 categories for all 15,235 species represented by ≥1 genome, and 22 PGPs across seven categories for 786 species represented by ≥10 genomes. These were integrated with phylogenetic, phenotypic and ecological data (Supplementary Table 3-6). Metadata and key process outputs, including whole-genome alignments and variation profiles, are available as community resources. We illustrate the utility of MAP across five distinct aspects.

### Quantitative characterization of prokaryotic species

First, MAP quantitatively characterizes prokaryotic species by ranking GPs and PGPs relative to all available species (**Fig. 2a,b**). For example, it confirms that the human pathogen *Helicobacter pylori* is highly recombinogenic^15^, with recombination-derived diversity (‘recombination coverage’, *Rec. c*^16^) in the top 1% and short-range (10 bp) linkage disequilibrium (LD, inversely correlated with recombination) in the bottom 1%. This quantitative characterization also revises prevailing views. For example, the widely distributed *E. coli*, naturally presumed to have a large effective population size (*N_e_*) and the correspondingly high diversity^13^, instead shows moderately high but not extreme *N_e_* (>71% species; estimated from *dN/dS*, a negative proxy for *N_e_* and selection efficacy in many microbial studies^17^) and intermediate genetic diversity (>39% species).

**Figure 2.**
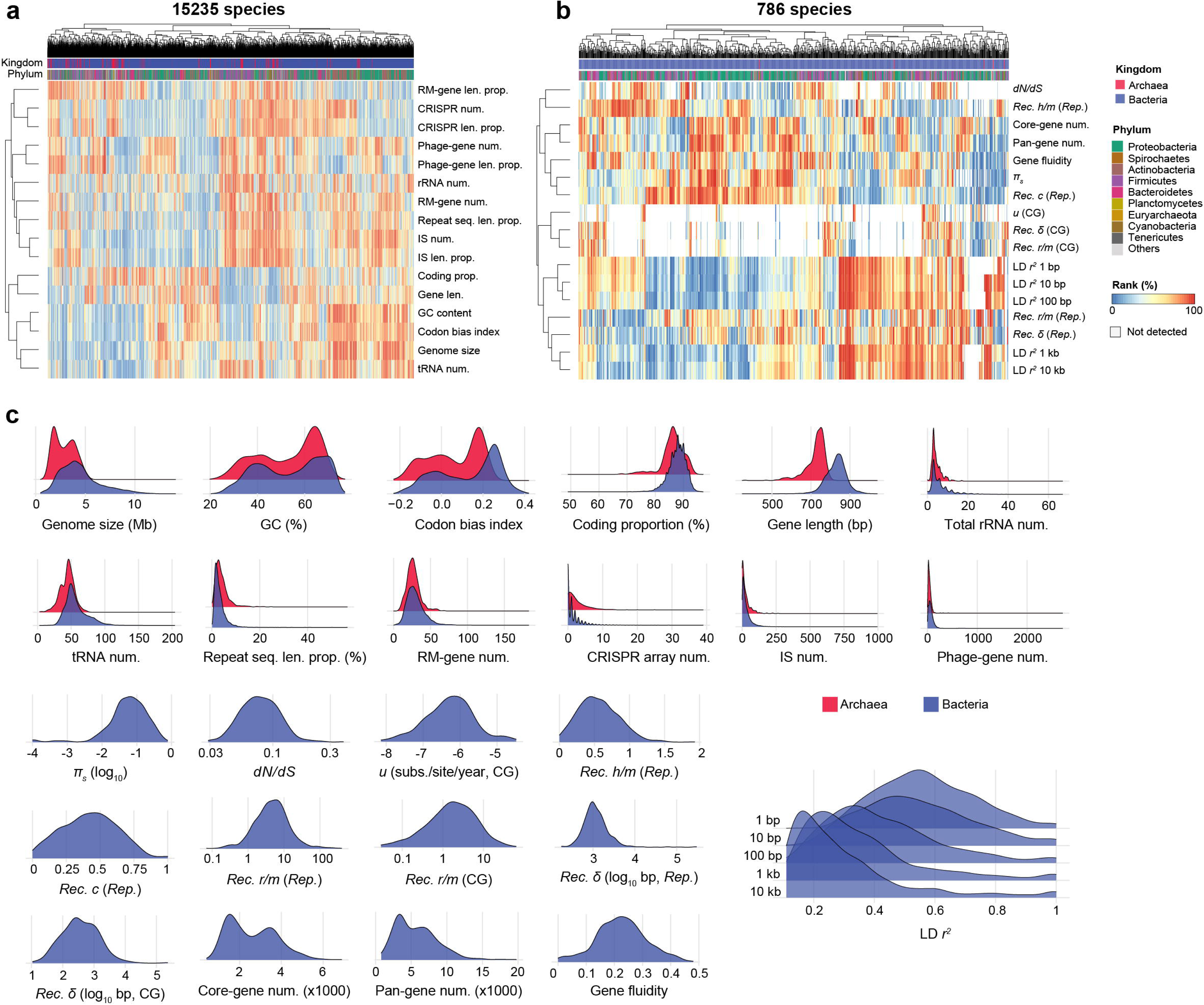
Overview of the MAP. **a,b)** Heatmaps showing the rank percentages of genomic parameters (GPs) and population genetic parameters (PGPs) for each species. Bars at the top indicate the kingdom and phylum, while heatmap colors represent rank percentages. **c)** Distribution of different GPs and PGPs. Red and blue represent archaea and bacteria, respectively. PGP is shown only for bacteria, as few archaeal species met the criterion for PGP estimation (≥10 representative genomes). *Rep.*: representative dataset; CG: clonal group dataset; num.: number; len.: length; prop.: proportion; *Rec.*: recombination-related parameters; *c*: recombination coverage; δ: recombination fragment size; *r/m*: relative effect of recombination to mutation; *u*: substitution rate; *h/m*: ratio of homoplasic and non-homoplasic alleles; *dN/dS*: ratio of non-synonymous divergence (*dN*) to synonymous divergence (*dS*); π*_s_*: synonymous genetic diversity. Source data are provided as a Source Data file.

### Defining range of prokaryotic diversity

Second, MAP delineates the range of genomic features across prokaryotes (**Fig. 2c**). For example, prokaryotes are generally small and densely coded^18^, with MAP revealing a median genome size of 4.0 Mb (interquartile range, IQR: 2.9–5.3 Mb) and coding proportion of 88% (IQR: 86–90%). Genomes larger than 7.3 Mb (top 10%), such as *Burkholderia cenocepacia* with multipartite chromosomes^19^, define the upper extremes.

This quantitative framework enables systematic identification of “extreme” species. For example, *H. pylori* carries an unusually high proportion of restriction-modification (RM) system genes (5.2% of the genome, top 1%), potentially reflecting their role in maintaining genome integrity while the species is naturally competent for foreign DNA uptake^20^. *Bordetella pertussis* has an exceptionally high proportion of insertion sequences (IS, 7.3% of the genome, top 1%), possibly providing alternative evolutionary routes via IS-mediated rearrangements^21^ in this otherwise extremely low-diversity species (bottom 1%). Other extremes, which seem to be unreported, include highly fluid genomes in gut-dominant *Bacteroides*, such as *B. uniformis* (fluidity =0.43, 43% of genes fluidic; top 1%) and *B. fragilis* (top 5%), likely due to frequent exposure to exogenous DNA^22^, and extremely low fluidity in intracellular pathogens, such as *Brucella melitensis* (bottom 1%, =0.01) and *B. abortus* (bottom 2%), likely reflecting their restricted lifestyles^23^.

### Overall patterns of prokaryotic genomes

Third, MAP reveals overall patterns of prokaryotic genomes (**Fig. 2c**). Archaea and bacteria show broadly similar GP distributions, except that archaeal genes are generally shorter. Most parameters follow roughly normal or right-skewed distributions. In contrast, GC content and codon bias index (CBI) are distinctly bimodal, with GC likely reflecting ancient adaptive legacies^24^, and CBI strongly influenced by GC^25^. Bacterial genome size approximates a normal distribution, whereas core- and pan-gene counts are bimodal. The distribution of core-genes likely reflects different outcomes of an evolutionary tradeoff but with the constraint that a minimum number of genes is needed for cellular function.

### Easy testing and refining evolutionary hypotheses

Fourth, MAP systematically quantified correlations among parameters (*r* for quantitative traits, η for categorical traits) and their relationships with phylogeny (Pagel’s λ), phenotype, and ecology (**Fig. 3a**). Although these correlations do not necessarily imply causal relationships, the resulting standardized correlation matrices provide an empirical resource for testing and refining evolutionary hypotheses.

**Figure 3.**
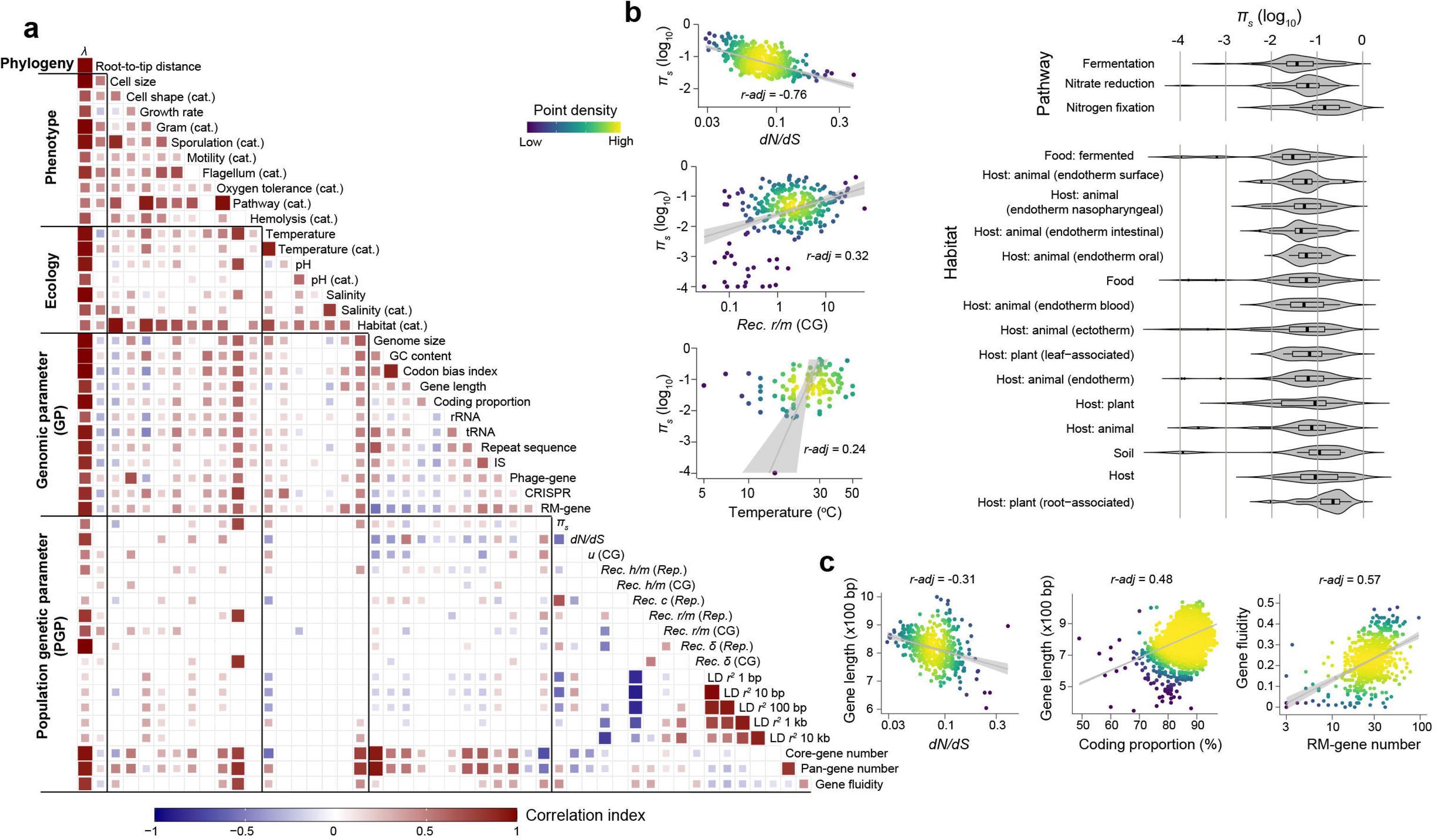
Correlation landscape of MAP. **a)** Correlation heatmap among various parameters. Phylogenetic signals were quantified using Pagel’s λ. cat.: categorical data. **b)** Correlations associated with genetic diversity (left) and genetic diversity across species from different sources (pathways or habitats, right). For categorical source variables, only categories with ≥10 species are shown. For box plots, the center line indicates the median, the box boundaries indicate the 25th and 75th percentiles. **c)** Underexplored correlations revealed by MAP. *r-adj* represents the correlation coefficient after adjustment for phylogenetic signal. Point colors represent point density. The grey line shows the fitted linear relationship, and the shaded area represents the 95% confidence interval. Source data are provided as a Source Data file.

### Determinants of genetic diversity

Our understanding of the determinants of genetic diversity remains limited. *N_e_*, mutation, and recombination have been proposed as major contributors^26^. While partly tested in eukaryotes, no equivalent dataset existed for prokaryotes because it is challenging to directly measure these factors. Two MAP parameters most closely related to these proposed contributors are *dN/dS* (*r-adj* = −0.76, inversely reflecting *N_e_*) and the recombination-to-mutation ratio (*r/m*, *r-adj* = 0.32), a proxy for recombination^27^. Both correlated with synonymous diversity π*_s_* in the expected direction, providing preliminary empirical support for their roles in shaping prokaryotic genetic diversity (**Fig. 3b**).

MAP also highlighted ecological associations with genetic diversity. Plant-root-associated species exhibited the highest π*_s_*, consistent with nutrient-rich conditions sustaining large populations, whereas fermented-food species showed the lowest, reflecting artificial selection and bottlenecks. Diversity also increased with temperature (*r-adj* = 0.24), consistent with previously reported latitudinal diversity gradients in both bacteria^28,29^ and eukaryotes^11^ (**Fig. 3b**).

### Underexplored correlations

MAP revealed previously overlooked patterns (**Fig. 3c**). Beyond confirming the established positive relationship between prokaryotic genome size and selection efficacy^17^, evidenced by the negative correlation between genome size and *dN/dS* (*r-adj* = −0.22, a negative proxy for selection efficacy), MAP further revealed an even stronger negative correlation between gene length and *dN/dS* (*r-adj* = −0.31). This pattern suggest that, analogous to genome size, stronger selection may favor longer genes. Gene length also correlated positively with coding proportion (*r-adj* = 0.48), possibly because regulatory non-coding regions remain relatively constant regardless of gene size.

MAP further uncovered an intriguing correlate of gene fluidity: RM-gene number correlated strongly with gene fluidity (*r-adj* = 0.57), implying selection for abundant RM systems under frequent mobile-element invasion, given their role in regulating DNA transfer^31^.

### Overview of prokaryotic genetic diversity

Finally, we illustrate how MAP can refine existing theoretical frameworks and generate new hypotheses. We focused on the relationship between LD and genetic diversity, summarized in a two-dimensional triangular plot, since short-range (10 bp) LD is intrinsically constrained to be higher than long-range (10 kb) LD (**Fig. 4a**). First, nearly the whole triangle is occupied, implying short- and long-range LD vary largely independently. Second, π*_s_*showed a strong negative correlation with short-range LD (*r-adj* = −0.58) but no significant correlation with long-range LD. Instead, for species with similar short-range LD, there is a pronounced positive correlation between long-range LD and π*_s_* (**Fig. 4a**), contrasting with the theoretical expectation that stronger genetic linkage should be associated with lower genetic diversity^26^. Together, these observations suggest that short- and long-range LD capture distinct aspects of evolutionary dynamics.

**Figure 4.**
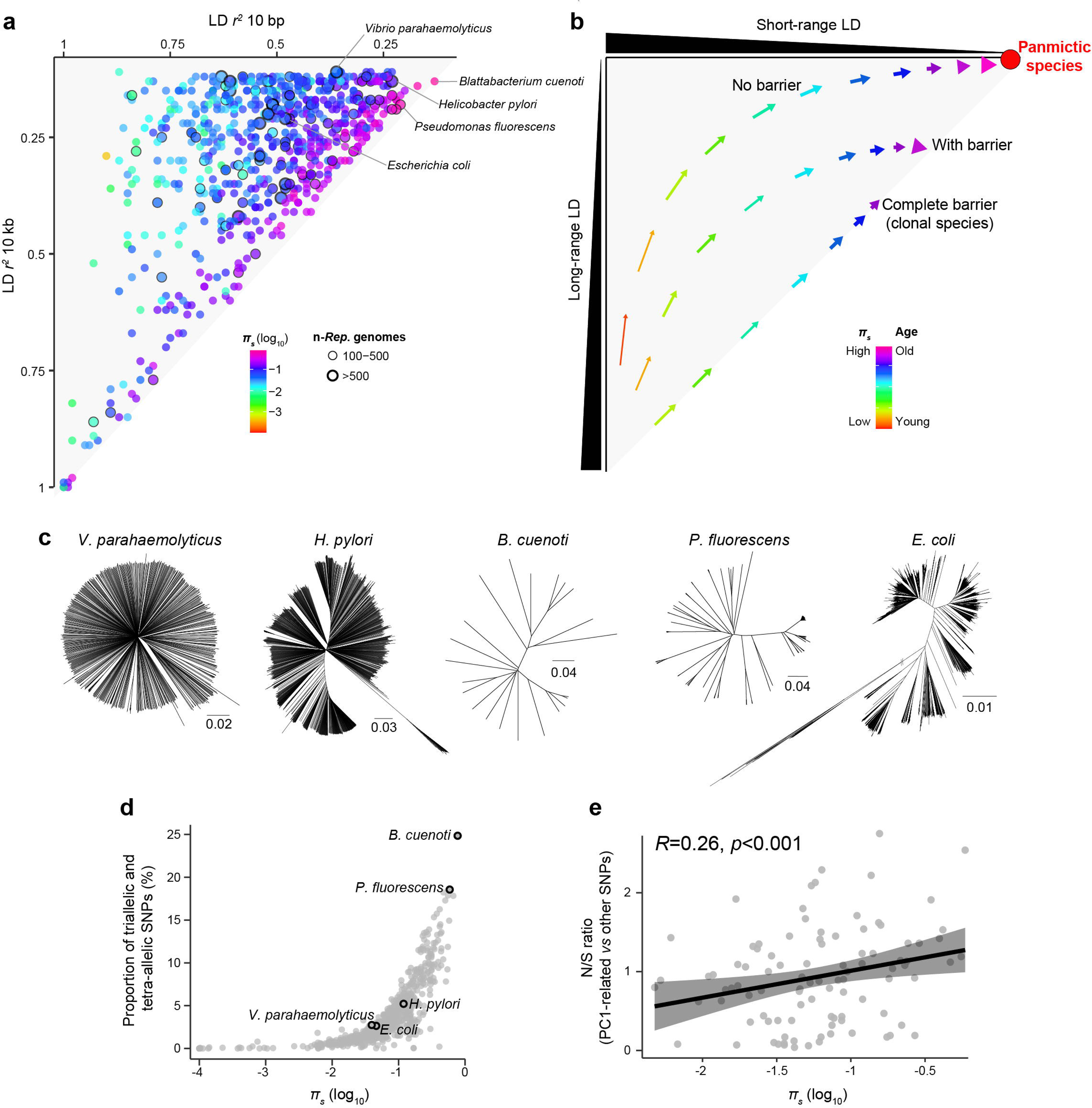
Overview of prokaryotic genetic diversity. **a)** Relationships between short-range (10 bp), long-range LD (10 kb) and genetic diversity. Point colors represent genetic diversity levels. Species with ≥100 representative genomes are outlined in black. **b)** Conceptual model illustrating the evolution of LD, genetic diversity, and population structure. The x- and y-axes represent the strengths of short- and long-range LD, respectively. Colored arrows depict evolutionary trajectories of species with or without genetic barriers, with colors indicating species ages reflected in genetic diversity. Panmictic species denote freely recombining and highly diverse species with minimal LD at all distances. **c)** Phylogenetic trees of five representative species. **d)** Correlation between genetic diversity and the proportion of triallelic and tetra-allelic SNPs. Five representative species are highlighted with black outlines. **e)** Correlation between genetic diversity and the N/S ratio, defined as the ratio of non-synonymous to synonymous sites for SNPs in the top 1% of loadings on PC1 (PC1-related SNPs) relative to the remaining SNPs. Pearson correlation was used to assess correlations, and the correlation coefficient (*R*) was calculated. The black line shows the fitted linear relationship, and the shaded area represents the 95% confidence interval. Source data are provided as a Source Data file.

These qualitative patterns can be interpreted within a conceptual framework defined by the joint effects of two variables: “species age”, reflected in genetic diversity, and barriers to gene flow (with non-recombining clonal species representing a complete barrier), reflected in long-range LD (**Fig. 4b**). In this framework, as species accumulate age (indicated by increased π*_s_*) in the absence of barriers, long-range LD decays rapidly, whereas short-range LD equilibrates more gradually, ultimately toward panmixia with high diversity and little LD at any distance (**Fig. 4b**, top arrows). By contrast, barriers to recombination slow the decay of long-range LD, raising its equilibrium level (**Fig. 4b**, middle and bottom arrows). Because LD decays more slowly in the presence of gene flow barriers, species positioned along a given vertical line in **Fig. 4a** tend to be “older” for higher long-range LD values (**Fig. 4b**), thereby explaining the positive association with π*_s_* along each vertical line.

Although simplified, for example because species-wide bottlenecks can reduce diversity and elevate LD, thereby making species appear “younger”, this two-dimensional framework provides a useful lens for interpreting species-specific LD and genetic diversity configurations.

The framework is consistent with several well-characterized species. A species with low barriers to genetic exchange is *Vibrio parahaemolyticus* (**Fig. 4a,c**), which exhibits low long-range LD but moderate diversity and correspondingly moderate short-range LD. Previous analyses suggested that its diversity remains moderate because population size expanded only recently, and if maintained over longer timescales, diversity would increase considerably, driving the species toward panmixia^32^.

A species with well-recognized barriers to genetic exchange is *E. coli* (**Fig. 4a,c**), where recombination between phylogroups is constrained by homology-dependent mechanisms^33^, resulting in higher long-range LD than *V. parahaemolyticus*. Another example is *H. pylori*, a species notorious for its exceptionally high rates of recombination during mixed infection^15^, which have led to very low short-range LD. However, its geographical structure, due to low rates of spread between human hosts as they dispersed out of Africa^34^, also leads to higher long-range LD than in *V. parahaemolyticus*.

Strikingly, although many species resemble *V. parahaemolyticus* in exhibiting low long-range LD, none approach complete panmixia. In our dataset, nearly all species display higher long-range LD than *V. parahaemolyticus* (**Fig. 4a**). Given that many bacteria occupy niches favorable to global dispersal and the maintenance of vast, stable populations over evolutionary timescales, there is therefore a paradoxical absence of panmictic species.

Indeed, the species with the lowest short-range LD, namely *Blattabacterium cuenoti* (**Fig. 4a**), is not panmictic. As an ancient intracellular cockroach endosymbiont, it reproduces strictly asexually and is therefore expected to lack recombination^35^. Consistent with this, we are unable to find unambiguous evidence of recombination, as reflected by its “flat” LD decay curve (Supplementary Fig. 5). Instead, *B. cuenoti* appears mutation-saturated, exhibiting an extreme excess of triallelic and tetra-allelic SNP sites compared to other species (**Fig. 4d**), indicative of genome-wide mutation saturation in an old and diverse species. Thus, its low LD likely arises from repeated mutation rather than recombination. No other species shows a comparable genome-wide mutation saturation signal in our dataset, and other diverse species, such as *Pseudomonas fluorescens*, show clear evidence of LD decay with distance (Supplementary Fig. 5). Although partial saturation at individual loci cannot be excluded, it is unlikely to substantially affect our genome-wide observation that panmictic species are rare.

### Role of epistatic selection in shaping population structure

One possible explanation for the paradoxical absence of true panmictic species is that species with low barriers to exchange systematically evolve stronger barriers as species age and accumulate genetic diversity. *V. parahaemolyticus* illustrates this process: in addition to weak geographic barriers generating oceanic gene pools^36^, further barriers arise from epistatic selection. Specifically, two major ecospecies are highly differentiated at ∼1% of the core genome, reflecting strong selection against gene flow in these regions^37,38^. If functional diversification expands over time, epistatic selection could progressively create barriers to exchange across larger genomic regions. *V. parahaemolyticus* would then not evolve into a panmictic species but instead come to resemble the structured, high-diversity species that we have observed.

We therefore propose that epistatic selection is one mechanism that could contribute to the absence of panmictic species, although it is unlikely to be the only one. Specifically, we hypothesize that panmictic bacteria are absent because, in species maintaining large *N_e_*, inevitable functional diversification driven by epistatic selection progressively generates long-range LD. This predicts stronger signatures of selection in the genetic regions responsible for population structure in “older” species with higher genetic diversity.

To evaluate whether this prediction is consistent with the data, we performed principal component analysis (PCA) for each species with >100 genomes and identified SNPs most strongly associated with the first principal component (top 1% by loading values; PC1-related SNPs), representing the genetic basis of major population structure^39^. As a proxy for selection, we quantified the enrichment of non-synonymous sites among these SNPs relative to the genomic background, using the N/S (non-synonymous to synonymous site) ratio.

Consistent with this prediction, enrichment of non-synonymous sites was positively correlated with π*_s_*, indicating that selection is more evident in highly diverse species (**Fig. 4e**). For example, *P. fluorescens*, which exhibits very high genetic diversity, showed strong enrichment of non-synonymous sites (N/S ratio = 2.5) and a highly structured phylogeny (**Fig. 4a,c**). Together, these results are consistent with our proposed framework in which epistatic selection progressively shapes population structure as species age and accumulate genetic diversity.

### Identification of convergent ecospecies

Although the proposed framework remains to be rigorously tested, it provides a practical strategy for identifying selection-driven population structures, such as ecospecies, which are characterized by fixed differences in a small fraction of the genome, but free genetic exchange elsewhere^37,38,40^. Guided by this framework, we searched for signatures of ecospecies in highly diverse species, asking whether any had progressed further along the road to functional differentiation than *V. parahaemolyticus*.

To this end, we examined species with higher diversity and many genomes (>100 genomes). Two such species are *Streptococcus mitis* and *S. oralis*. The PCA loading distribution for both resembled that seen for previous ecospecies^37,38,40^, showing strong differentiation in specific genomic regions together with high gene flow elsewhere (**Fig. 5a,b**). Furthermore, there is strong enrichment of genes of particular categories with extreme PC1 loadings. For both species, COG categories D (cell cycle control, cell division, chromosome partitioning) and M (cell wall/membrane/envelope biogenesis), and KEGG category ko03036 (chromosome and associated proteins) were involved, suggesting functional divergence in cell division and biogenesis. The convergence of the differentiation signal is further supported by a correlation of 0.41 between PC1 gene loadings in the two species, together with substantial overlap among genes with extreme PC1 loadings in the D, M and ko03036 categories (**Fig. 5c**).

**Figure 5.**
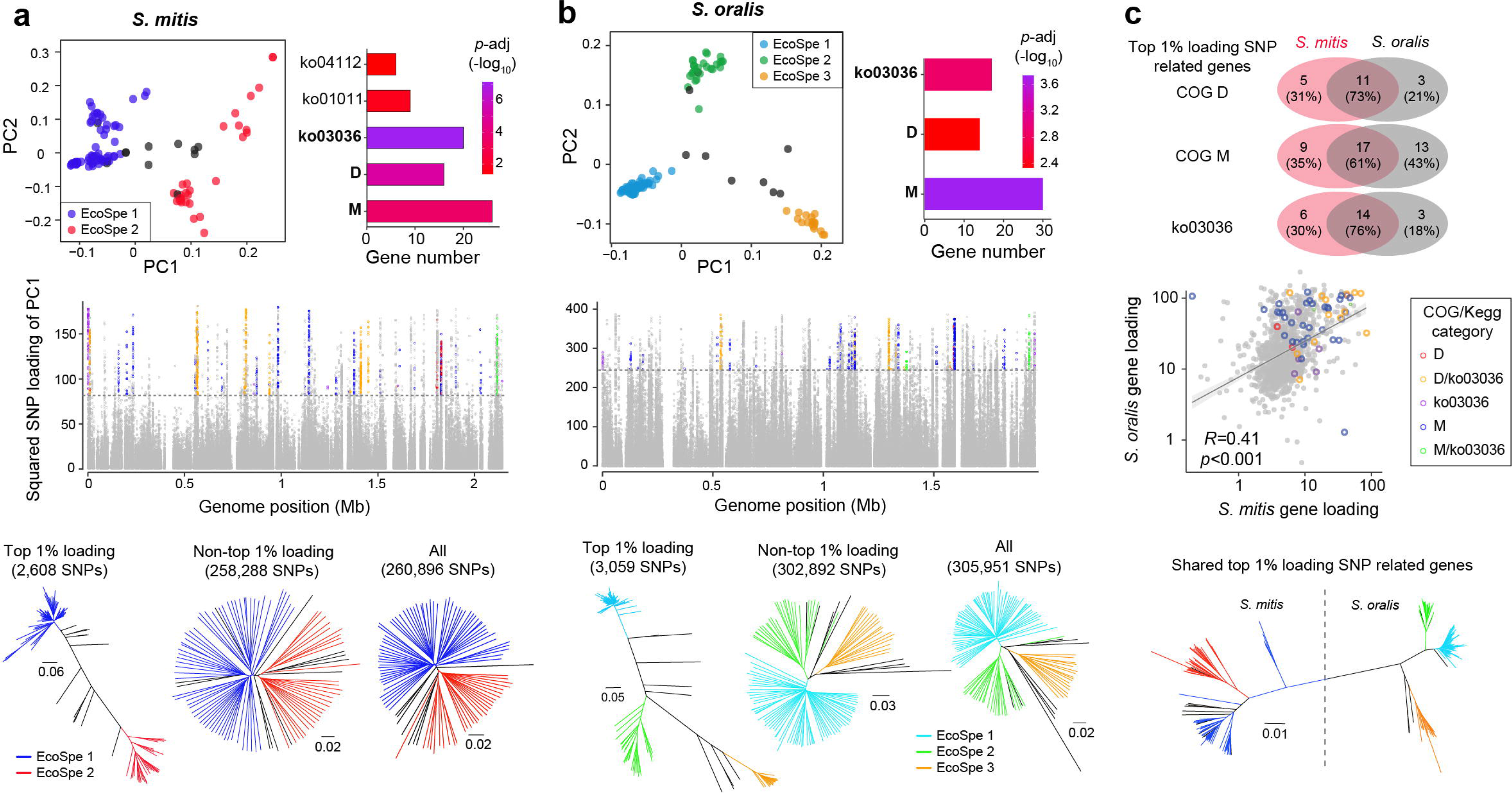
Convergent ecospecies. **a,b)** Top: Principal component analysis (left; point colors indicate inferred ecospecies) and functional enrichment of genes associated with PC1-related SNPs (top 1% by loading values; right) in *Streptococcus mitis* and *S. oralis*. Middle: Genome-wide distribution of SNP loadings on PC1. Point colors represent COG or KEGG categories. Dashed lines indicate the top 1% loading cutoff. Bottom: Phylogenetic trees based on PC1-related SNPs, the remaining SNPs and all SNPs. Branch colors indicate inferred ecospecies. **c)** Top: Shared homologous genes associated with PC1-related SNPs between the two species. Middle: Correlation of gene loading (mean SNP loading per gene) between species. Point colors represent COG or KEGG categories. Bottom: Phylogenetic trees based on protein sequences of shared genes associated with PC1-related SNPs. Pearson correlation was used to assess correlations, and the correlation coefficient (*R*) was calculated. The grey line shows the fitted linear relationship, and the shaded area represents the 95% confidence interval. Source data are provided as a Source Data file.

To characterize these putative ecospecies, we constructed phylogenetic trees based on SNPs with extreme loadings. We found two putative ecospecies for *S. mitis* and three for *S. oralis*, which also correspond to distinct clusters in strain PCA loadings (**Fig. 5a,b**). To determine whether they had evolved independently, we constructed a joint tree for the shared high loading genes, which implies independent origin of ecospecies in the two species (**Fig. 5c**). Finally, we constructed phylogenetic trees based on remaining SNPs and all SNPs to examine the pattern of differentiation in the rest of the genome. Distinct ecospecies clusters are visible, implying modest genome-wide differentiation, which contrasts with the absence of differentiation in most of the genome seen in *V. parahaemolyticus* and *H. pylori*^37,40^. This result is consistent with the *Streptococcus* ecospecies being older and further along the path of functional differentiation and helps to explain the higher long-range LD seen in these two species.

## Discussion

Exploring and understanding the diversity of life requires a framework akin to a geographical map. For prokaryotes, classical resources such as *Bergey’s Manual* have long served as the initial “map”. MAP extends this foundation with a genome-based atlas, capturing rich genomic and population genetic features across prokaryotic species. It is accessible online, enabling straightforward exploration and use.

MAP serves both focused and broad research needs. For researchers interested in particular microbes, it provides concise summaries of genomic, population genetic, phenotypic, and ecological features. Beyond single species, it facilitates straightforward comparisons across all available prokaryotes and serves as a resource for macrogenomic and population genetic studies, including both raw and curated non-redundant datasets. The current MAP is primarily based on genomes from isolated strains, and future versions will incorporate metagenome-assembled genomes to expand species coverage and genomic diversity.

Our analysis of MAP has generated a number of interesting observations that deserve further study. For example, while bacterial genome size approximates a normal distribution, core- and pan-gene counts are unexpectedly bimodal. Gut-dominant *Bacteroides*, such as *B. uniformis* and *B. fragilis,* have exceptionally high genome fluidity. Moreover, we confirm that selection favors bigger genomes^17^, and also find a negative correlation between gene length and *dN/dS*, suggesting that selection also favors long genes.

Our analyses and resources are available based on two taxonomic classifications, namely NCBI and Genome Taxonomy Database (GTDB)^41,42^. The problem of bacterial species definitions remains unsolved and large databases inevitably contain errors^43,44^. NCBI is based on user inputted classifications, while GTDB uses an automatic algorithm, based on a universal average nucleotide identity cutoff of 0.95. While GTDB has advantages of consistency and quality control over NCBI, it often splits taxonomically diverse named species into multiple putative species (Supplementary Fig. 6). In many cases, these species do not correspond to coherent evolutionary units. In particular, highly diverse species such as *P. fluorescens* have very low short-range LD, indicating that gene flow is effective in distributing diversity within the species, and accordingly they are among the best examples of biological species, despite being split into >10 species by GTDB. Therefore, GTDB is not especially well-suited for macrogenomic analysis, especially for highly diverse species, and we presented our main results based on NCBI classification. We nevertheless provide an alternative version of parameter calculation and correlation analysis results based on GTDB (Supplementary Tables 7-9), and show a strong correlation between parameter relationships derived from two taxonomies (*r* = 0.96, Supplementary Fig. 7), indicating that the great majority of our results are robust to species definitions.

The correlation matrix integrating genomic, population genetic, phenotypic, and ecological parameters provides a resource-level overview that facilitates hypothesis generation and testing, thereby helping to refine existing theoretical frameworks. These correlations, however, do not necessarily imply causal relationships and may include false positives arising from multiple testing or sensitivity to outliers; they should therefore be interpreted with caution. Nonetheless, they serve as a valuable starting point for hypothesis-driven and statistically rigorous follow-up analyses.

As an illustration, motivated by the observed lack of correlation between short- and long-range genetic linkage, we propose and provide evidence that these two linkage scales capture distinct aspects of evolutionary dynamics and independently shape genetic diversity. Classical population genetic theory predicts a negative correlation between genetic diversity and genetic linkage level^26^. We indeed find that short-range LD is strongly inversely correlated with genetic diversity. However, there is a pronounced positive correlation in MAP between long-range LD and π*_s_* for species exhibiting similar levels of short-range LD (**Fig. 4a**). We propose that this pattern can be interpreted by considering the joint effects of “species age” and barriers to gene flow. Because long-range LD principally reflects barriers to gene flow, it decays more slowly across all genetic distances (**Fig. 4b**). Consequently, among species with similar short-range LD values, those with higher long-range LD tend to be older and more diverse. Although simplified, this conceptual framework is consistent with several well-characterized species and provides a useful basis for generating testable evolutionary hypotheses.

There is one zone in the triangle that is surprisingly empty, corresponding to panmictic species with low LD at all distances (**Fig. 4a**). Many bacteria occupy niches favorable to global dispersal and the maintenance of vast, stable populations over evolutionary timescales and this should generate panmictic, i.e., freely mixed species^32^. Our resolution of this paradox is that as within-species diversity increases, functional diversification driven by epistatic selection progressively generates barriers to gene flow. Consistent with this, we identified stronger signatures of selection structuring diversity in more genetically diverse species (**Fig. 4e**). Thus, while there may be species that start on the journey to panmixia, with large population sizes, weak geographic barriers and free recombination (as exemplified by *V. parahaemolyticus*), they may progressively acquire barriers to gene flow due to epistatic selection as they age, and end up with intermediate long-range LD. Together, these observations are consistent with epistatic selection as one mechanism that could contribute to this process, although it is unlikely to be the only one. Other processes, including geographic isolation and intrinsic barriers to genetic exchange^33,34^, are also likely to play important roles.

An important implication of our findings is that we can gain insight into the ecological pressures acting on genetically diverse prokaryotic species, simply by characterizing how their diversity is structured. One example of structure generated by selection is bacterial ecospecies^37,38,40^. We have previously described the first two bacterial ecospecies in *V. parahaemolyticus* and *H. pylori.* In *V. parahaemolyticus* the structure reflects diversification at motility-related genes^37,38^, while in *H. pylori* differentiation has occurred in genes related to immune escape and nutrient uptake^40^. Both species have moderate diversity and ecospecies structure is restricted to a small fraction of the genome, representing an early stage of diversification.

Guided by this framework, MAP provides a practical strategy for identifying additional candidate ecospecies. Here we have identified two additional species with ecospecies-like structure, namely *S. mitis* and *S. oralis*. These two sister taxa both inhabit the mouths of humans and other animals and there is a substantial convergence in the genetic basis of the ecospecies structure, with the same cell division and biogenesis associated genome categories and genes involved. These suggest similar ecological driving forces and/or adaptive potentials in both species, which might reflect the diverse niches within complex multispecies biofilms in the oral cavity, that could be exploited by evolving cell shape and cell wall properties. The ecospecies appear to be older than those identified previously and show evidence for some level of differentiation genome-wide, suggesting a later stage of diversification and is likely to be a precursor to full speciation. Together, these findings demonstrate how MAP can generate evolutionary hypotheses and facilitate the discovery of ecospecies.

Extrapolating from early to late stage ecospecies divergence and beyond, we can begin to reconstruct the life-cycle of many prokaryotic species. However, ecospecies is only one type of structure caused by selection. We have previously hypothesized the existence of bacterial ecoclines, which exhibit continuous variation driven by a quantitative trait^39^ and the genetic structure in the most diverse species such as *P. fluorescens* resists any immediate classification. Characterizing the full range of prokaryotic genetic structures should provide a unique new window into the forces shaping the most diverse and adaptable organisms on our planet.

## Methods

### Genome datasets

#### Deduplication and quality control

We retrieved all assembled genomes (n=317,542) included in the Genome Taxonomy Database (GTDB; https://gtdb.ecogenomic.org, release 207)^41,42^. We performed genome deduplication based on BioSample accessions, retaining only the genome with the highest N50 value for each BioSample. Only high-quality genomes (n=268,463) with estimated completeness ≥90% and contamination ≤5%, as assessed by CheckM v1.1.3^45^, were retained for subsequent analyses.

The high-quality genomes were assigned to species based on both National Center for Biotechnology Information (NCBI) taxonomy and GTDB classifications. For each species, the earliest sequenced complete genome, or alternatively the genome with the highest N50 value (if no complete genome was available), was designated as the reference genome. All genomes of a species were aligned to the corresponding reference genome using MUMmer v3.23^46^ to generate whole-genome alignments, as previously described^36^, and only genomes with ≥60% alignment coverage against the reference were retained for subsequent analyses.

After deduplication and quality control, a total of 266,627 high-quality genomes were retained for downstream analyses, and their genome quality metrics and taxonomic classifications are provided in Supplementary Table 1. The whole-genome alignments for each species are publicly available from the Zenodo database (https://zenodo.org/records/17430471).

#### Representative datasets and clonal groups

While robust estimation of population genetic parameters requires randomly sampled representative datasets, biased repeat sampling (*e.g*., strains from disease outbreaks) is prevalent in public genomic datasets, especially for pathogenic species. To reduce the effects of repeat sampling, we generated representative genome datasets as previously described^47^. Briefly, for each species, core-genome (regions present in >95% strains) single-nucleotide polymorphisms (SNPs) were identified using SNP-sites v2.5.1^48^. SNPs located in repetitive regions, as identified by TRF v4.07b^49^ or BLASTN self-alignment, were excluded from subsequent analyses. Pairwise SNP distances between strains were calculated using snp-dists v0.8.2 (https://github.com/tseemann/snp-dists). Genomes of a species were grouped into complete-linkage clusters (CLCs) based on pairwise SNP distances using the R package ape v5.7.1^50^. For the representative dataset of each species, we used a “relative” threshold, defined as 1% of the median pairwise SNP distance, as the cutoff, and selected a single genome with the highest N50 from each CLC or from singleton genomes as representative genomes.

Estimation of substitution rates and selected recombination parameters rely on clonal groups (CGs) consisting of closely related genomes. We used an “absolute” SNP-distance threshold of 5,000 SNPs (corresponding to the total SNP cutoff per branch for reliable recombination detection^51^) as the cutoff to identify CLCs corresponding to CGs. The assignments of the representative and CG datasets are provided in Supplementary Table 1.

### Genomic parameters

We calculated genomic parameters (GPs) across 12 categories for all representative genomes, using the median value across representative genomes to represent each species (Supplementary Table 2 and Table 4). The variability of each parameter, quantified as the standard deviation (SD)-to-median ratio, is shown in Supplementary Fig. 2. Some GP estimates exhibited relatively high variability (SD/median ratio >0.5), which we interpreted as indicative of substantial within-species heterogeneity among strains. Rather than excluding these parameters, we retained them and used the median value as a representative measure, in order to maximize comparability across species.

Specifically, genome size and GC content were calculated using Quast v5.0.2^52^. We re-annotated representative genomes using Prokka v1.13^53^ with default settings, except for the “gcode” (genetic code) option. Genetic codes for each species were obtained from the NCBI taxonomy database. For species included in MAP, we found that all species use translation table 11 (Bacterial, Archaeal, and Plant Plastid Code), except for species classified under NCBI taxonomy “Mycoplasmatales and Entomoplasmatales” (153 species) or GTDB taxonomy “Mycoplasmatales” (198 species), which use translation table 4. Gene number and coding proportion were extracted or calculated based on annotation results. Codon bias parameters (CBI, Fop, ENc) were estimated using codonW v1.3 (https://codonw.sourceforge.net/) based on coding sequences. The number of tRNA and rRNA genes was extracted from Prokka annotations.

The standard protein sequences of restriction-modification (RM) system genes were retrieved from REBASE^54^ and searched against annotated protein sequences to identify RM genes using BLASTp; hits with >50% coverage were considered present as previously described^36^. Repetitive regions of genomes were identified using TRF v4.07b^49^ and BLASTN self-alignment as previously described^36^. CRISPR arrays were identified using CRISPRidentify v1.1.0^55^. Phage sequences were identified using PhiSpy v3.7.8^56^. Insertion sequence (IS) elements were identified using ISEScan v1.7.2.3^57^. Default parameters were used for all analyses unless otherwise specified.

### Population genetic parameters

We calculated seven categories of population genetic parameters (PGPs) for species with ≥10 representative genomes and for CGs with ≥10 genomes (Supplementary Table 5).

#### Random sampling procedure

To reduce potential biases arising from unequal numbers of representative genomes, five categories of parameters (π*_s_, dN/dS,* LD *r*^2^*, h/m,* core-/pan-genome and gene fluidity) were estimated using random sampling of 10 representative genomes. Sampling was repeated 10 times independently.

For *dN/dS* and LD *r²*, which require specific thresholds for reliable estimation (see below), some replicates failed to yield results. Only estimates with ≥5 successful replicates were retained for downstream analyses. To further ensure robustness, we excluded cases with unstable outcomes, defined as those with a SD/median ratio >0.5 across replicates. The final estimate for each species was taken as the median across retained replicates. The variability (SD/median ratio) of each parameter is shown in Supplementary Fig. 2.

#### π_s_ and dN/dS

For each species, coding sequence alignments of representative genomes were extracted from whole-genome alignments. Pairwise non-synonymous divergence (*dN*), synonymous divergence (*dS*), and *dN/dS* were calculated with PAML v4.10.0^58^ using the yn00 algorithm. The mean pairwise *dS* was used to represent the synonymous nucleotide diversity (π*_s_*). For *dN/dS* estimation, to reduce the impact of heterogeneous divergence levels across species while maximizing the number of species that could be compared, we retained only genome pairs with *dS* values within 0.03–0.13, corresponding to the interquartile range (IQR) across all species. For random sampling analyses, only *dN/dS* estimates based on >10 strain pairs were retained.

#### Linkage disequilibrium

Linkage disequilibrium (LD) correlation coefficient between two loci (*r^2^*) at distances of 1 bp, 10 bp, 100 bp, 1 kb, and 10 kb was calculated using Haploview v4.2^59^. For each replicate, estimates were retained only when >1,000 locus pairs were available at the given distance.

#### h/m

We calculated the ratio of homoplasic and non-homoplasic alleles (*h/m*) using ConSpeciFix v1.3.0^60^ based on randomly sampled and all representative genomes.

#### Core- and pan-genome and gene fluidity

For each species, the number of core- and pan-genes was calculated using Panaroo v1.2.8^61^ based on re-annotated results. Gene fluidity was measured by the mean pairwise ratio between the number of genes not shared and the total number of genes.

#### Recombination estimation based on all representative genomes

Five recombination-related parameters were estimated from all representative genomes using mcorr^16^ with default settings. These included mutational divergence (θ_pool_), recombinational divergence (φ_pool_), mean fragment size (δ), fraction of the sample diversity resulting from recombination (*c*), and the relative effect of recombination to mutation (*r*/*m* = φ_pool_ ⁄θ_pool_).

#### Recombination level and substitution rate of clonal group (CG)

For each CG, all available genomes were used to estimate recombination-related parameters and substitution rates. The variability (SD/median ratio) of each parameter across CGs within a species is shown in Supplementary Fig. 2. We interpreted high variability (SD/median ratio >0.5) as reflecting substantial within-species heterogeneity among CGs. Rather than discarding such cases, we retained them and used the median value as the representative estimate for each species, thereby maximizing comparability across species.

Four recombination-related parameters were estimated for each CG using ClonalFrameML v1.12^27^: the relative rate of recombination to mutation (*R/*θ), mean divergence between donor and recipient (ν), δ and *r*/*m*. In addition, *h/m* was estimated using ConSpeciFix v1.3.0^60^.

For each CG of a species, we reconstructed maximum-likelihood (ML) phylogenetic trees using FastTree v 2.1.10^62^ based on non-repetitive and non-recombined SNPs as previously described^63^. The isolation year of strains was extracted from the corresponding BioSample information. We used the ML tree and isolation year information to calculate the substitution rate with BactDating v1.1.0^64^ as previously described^47^. Only substitution rate estimates (number of substitutions per site per year) with effective sample sizes (ESS) exceeding 100 were retained, ensuring robust temporal signal and reliable estimation.

### Phenotype and ecology data

The phenotype and ecology data were extracted from three sources: Madin *et al*^65^, Secaira-Morocho *et al*^66^, and the BacDive database (up to August 2025)^67^. Data from the first two sources were available at the species level. For species with multiple strain entries, quantitative traits were summarized using the mean or median, while categorical traits were assigned according to the majority state (term occurring in >50% of strains).

BacDive provides strain-level data, which were aggregated to the species level using the same approach: the median for quantitative traits and the majority state for categorical traits. When no single state exceeded a 50% frequency, the categorical trait was recorded as “not available (NA)”. For BacDive-derived traits, within-species variability (SD/median ratio across strains) for quantitative traits and the frequencies of majority states across species for categorical traits are shown in Supplementary Fig. 2. Traits with high variability were retained to maximize comparability across species.

We further combined data from all three sources by taking the median of quantitative traits and the majority state across sources for categorical traits. The resulting species-level data from each individual source, as well as the combined dataset, are presented in Supplementary Table 3.

### Correlation analysis

Correlation analyses were performed using the Python packages pingouin and scipy.stats. All pairwise combinations of parameters were evaluated for parameter pairs containing more than 100 observations. For continuous quantitative parameters, Spearman correlation was used throughout to minimize sensitivity to outliers and differences in scale. We further compared Spearman, Kendall, and Pearson correlations to assess the robustness of the observed associations, which showed high concordance across methods (Supplementary Fig. 8). Associations between quantitative and categorical parameters were quantified using the eta coefficient (η), whereas associations between two categorical parameters were evaluated using Cramér’s coefficient (*V*).

The *P* values from all tested parameter pairs were adjusted for multiple testing using the Benjamini–Hochberg false discovery rate (FDR) procedure implemented in the Python package statsmodels using the multipletests function (method = ‘fdr_bh’). Associations with an FDR-adjusted *P* < 0.01 and *|r*|, η, or *V* > 0.1 were considered significant (Supplementary Table 6).

The phylogenetic signal of all parameters, including GPs, PGPs, and phenotypic and ecological parameters, was quantified using Pagel’s λ and estimated with the R package geiger v2.0.11^50^. Specifically, the fitContinuous function was applied with the model set to “lambda”, using phylogenetic trees based on universally distributed marker genes (archaea: 53 genes; bacteria: 120 genes) as implemented in GTDB^41,42^, together with the corresponding parameter values as input. Categorical traits were converted into continuous numerical values ordered from low to high, such that identical categorical states were represented by the same numerical value.

Phylogenetically corrected correlations for the specific parameter pairs shown in Fig. 3 were calculated using the R package phylolm v2.6.2^68^. The phylolm function was applied with the model set to “lambda”, incorporating the same GTDB-based phylogenetic trees and corresponding parameter values as input, with the regression formula specified as *parameter A ∼ parameter B*. Three parameters (π*_s_, dN/dS*, and *r/m*) were log10-transformed before phylogenetic correction because they showed skewed distributions in their original scales. The other five parameters (temperature, gene length, coding proportion, RM-gene number and gene fluidity) were analyzed using their original values.

### Principal component analysis (PCA)

PCA was performed using PLINK v1.9^69^ based on core-SNPs with a minor allele frequency (MAF) threshold of 2%. The SNP loading for each principal component (PC) was calculated as described in Liu *et al*.^39^, reflecting the correlation between SNPs and PCs. Briefly, strain loadings were derived by multiplying the eigenvectors by the square root of the eigenvalues corresponding to each PC. SNP loadings were then computed by matrix multiplication between strain loadings and the SNP matrix. SNPs in the top 1% of loading values for the first principal component were defined as PC1-related SNPs, representing the genetic basis of major population structure, whereas the remaining SNPs were treated as genomic background.

### SNP annotation and gene enrichment analysis

Synonymous and non-synonymous SNPs were annotated using SnpEff v4.3t^70^, based on the re-annotated reference genome. Genes associated with PC1-related SNPs were extracted for subsequent analysis. Functional enrichment was performed using the R package ClusterProfiler v4.2.2^71^.

### Between-species homologous gene analysis

Homologous genes between *Streptococcus mitis* and *S. oralis* were identified using BLASTp based on annotated protein sequences, with a minimum identity and coverage of 50%. Protein sequences of homologous genes associated with PC1-related SNPs were used for phylogenetic tree construction using FastTree v2.1.10^62^.

## Data availability

All genome sequences are available from the Genome Taxonomy Database (GTDB; https://gtdb.ecogenomic.org, release 207). Essential data required for replication, including whole-genome alignments, SNP matrices, curated metadata, and calculated results, are available at www.genomap.cn and Zenodo (https://zenodo.org/records/21357528), and are also provided in Supplementary Tables. Linux command-line scripts used to calculate genomic and population genetic parameters are provided in the Supplementary Text. Source Data are provided with this paper.

## Acknowledgements

This work was supported by National Key Research and Development Program of China (No. 2022YFC2304700 to F.D.), National Natural Science Foundation of China (No. 32270003 and No. 32000008 to C.Y., No.32350710791 and No. 32170640 to D.F.), Youth Innovation Promotion Association, Chinese Academy of Sciences (No. 2022278 to C.Y.) and Shanghai Rising-Star Program (No. 23QA1410500 to C.Y.).

## Author contributions

C.Y., D.F., R.Y., and Y.C. initiated and coordinated the study. C.Y., N.W., H.H. and W.X. contributed to data collection and performed bioinformatics analysis. F.J. and S.T. contributed to website construction and visualization. All authors contributed to interpretation of the data. C.Y. and D.F. wrote the first draft of the paper and D.X., R.Y., and Y.C. reviewed and revised the paper. All authors read and approved the final manuscript.

