## Supplementary Material for "Macrogenetic atlas of prokaryotes reveals selection-driven structures"

### Supplementary materials

**Supplementary Table 1.** High-quality genomes with species assignment and classification in the Representative and Clonal Group datasets.

**Supplementary Table 2.** Metadata and genomic parameter values for all high-quality Representative genomes.

**Supplementary Table 3.** Phenotypic and ecological data integrated from three publicly available data sources.

**Supplementary Table 4.** Species-level genomic parameter ranks and values for 15,235 species (NCBI taxonomy). Each table entry shows the rank (rank/total species) and the median±SD.

**Supplementary Table 5.** Species-level population genetic parameter ranks and values for 786 species (NCBI taxonomy). Each table entry shows the rank (rank/total species) and the median±SD.

**Supplementary Table 6.** Correlation matrices among genomic, population genetic, phylogenetic, phenotypic, and ecological parameters based on the NCBI taxonomy. a) Correlation matrices for all individual parameters. b) Correlation matrices between parameter categories, where each category–category correlation is represented by the maximum correlation coefficient among all parameter pairs within the two categories.

**Supplementary Table 7.** Species-level genomic parameter ranks and values for 42,469 species (GTDB taxonomy). Each table entry shows the rank (rank/total species) and the median±SD.

**Supplementary Table 8.** Species-level population genetic parameter ranks and values for 1411 species (GTDB taxonomy). Each table entry shows the rank (rank/total species) and the median±SD.

**Supplementary Table 9.** Correlation matrices among genomic and population genetic parameters based on the GTDB taxonomy.

**Supplementary Figure 1.** Distribution of representative genomes (a), complete linkage clusters (b), and clonal groups (c) per species. GTDB taxonomy is shown on the left, NCBI taxonomy on the right.

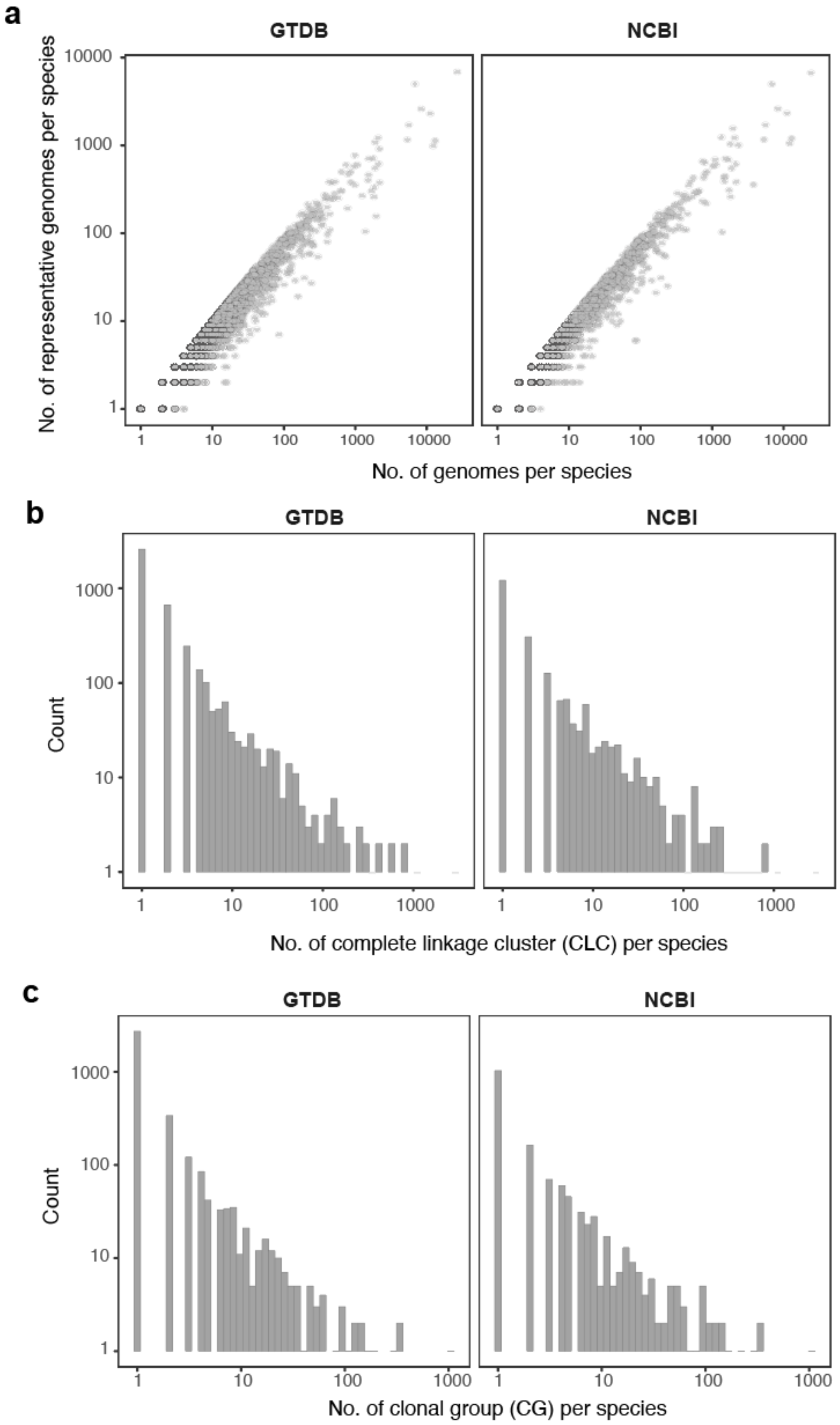

**Supplementary Figure 2.** Variability of genomic and population genetic parameters (NCBI taxonomy; a,b) and phenotypic and ecological parameters (BactDive database; c,d). Histograms show the distribution of each parameter across all species. For continuous data, variability for each species was calculated as the ratio of the standard deviation (SD) to the median across all representative genomes (a), random sampling replicates or distinct clonal groups within a species (b), or all strains with available data (c). For categorical data, the proportion of strains in each major category is shown for each species (d).

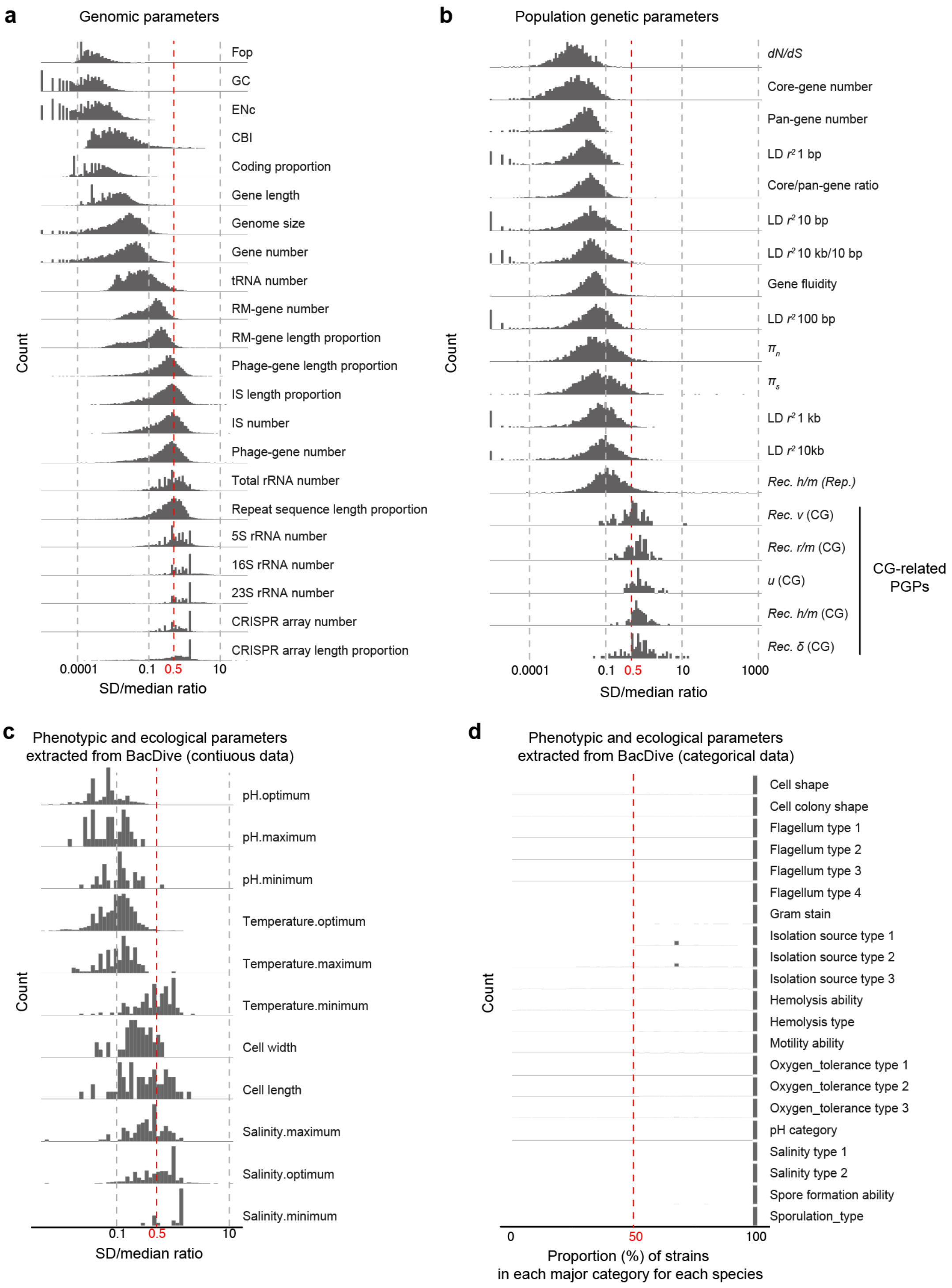

**Supplementary Figure 3.** Correlations between population genetic parameters (NCBI taxonomy) estimated from random sampling and those based on all representative genomes. *R*: correlation coefficient.

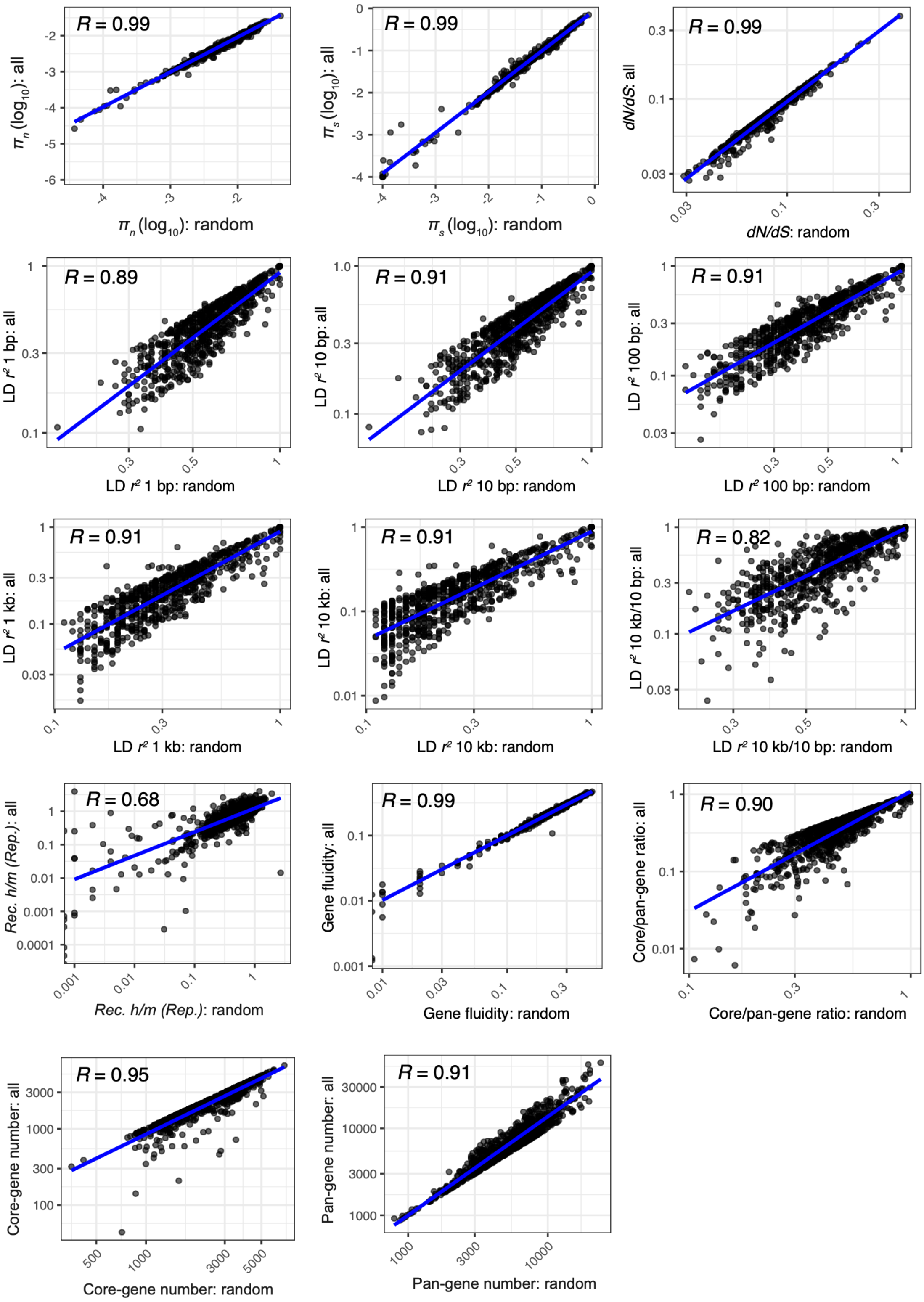

**Supplementary Figure 4.** Correlations between population genetic parameters (NCBI taxonomy) estimated from random sampling and number of representative genomes. *R*: correlation coefficient.

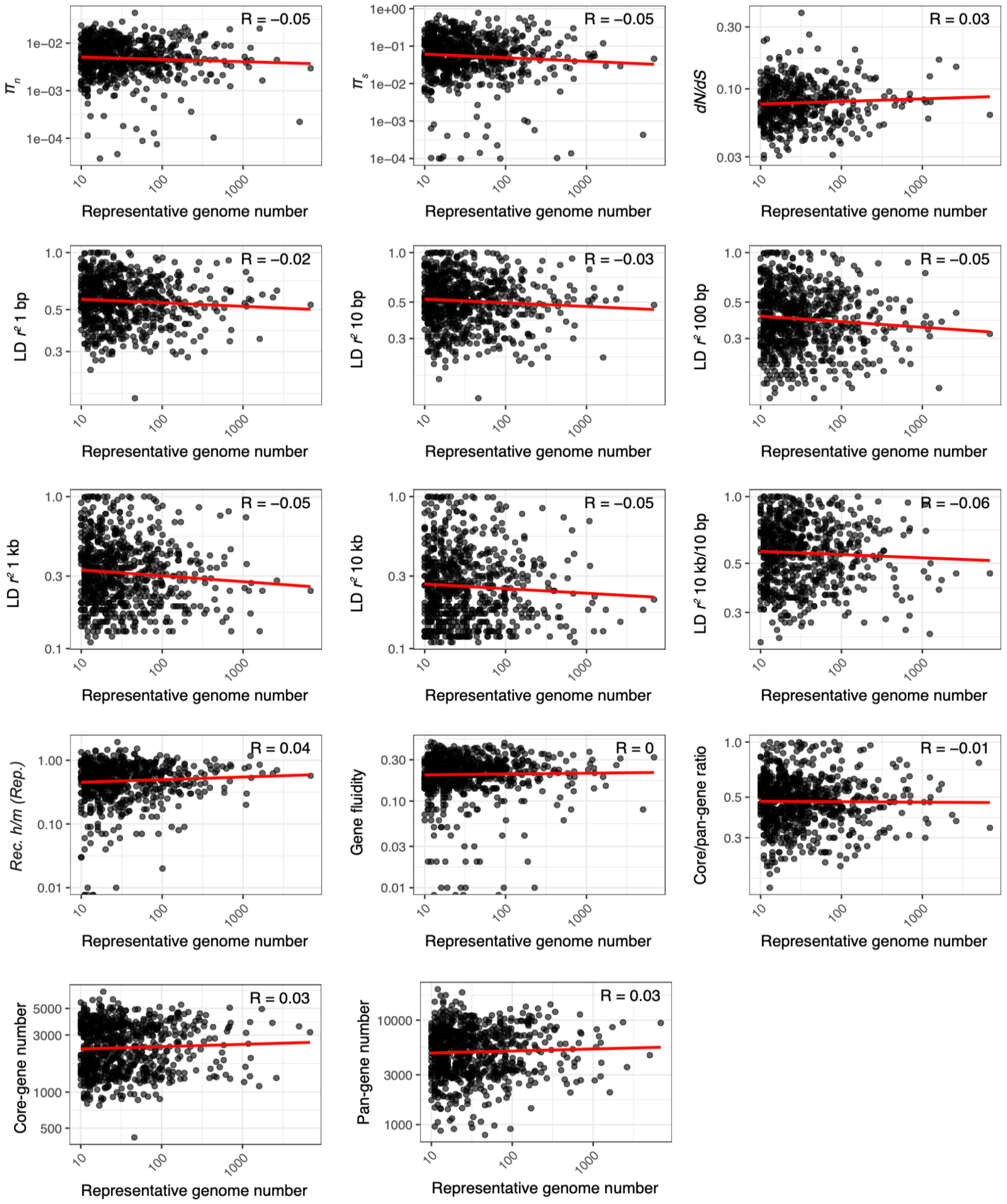

**Supplementary Figure 5.** LD decay curves of five representative species shown in Fig. 3a. The x-axis represents genetic distance, and the y-axis represents the linkage disequilibrium (LD) correlation coefficient between two loci ($r^2$). Data represent the median values from 10 replicates based on 10 randomly selected genomes.

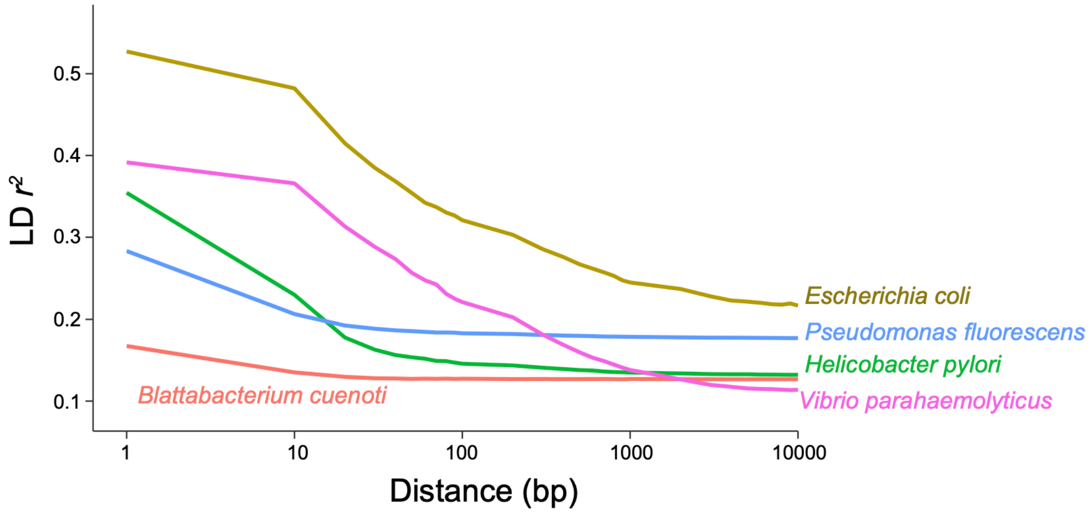

**Supplementary Figure 6.** Phylogenetic trees of the seven species shown in Figures 3 and 4 annotated with GTDB taxonomies.

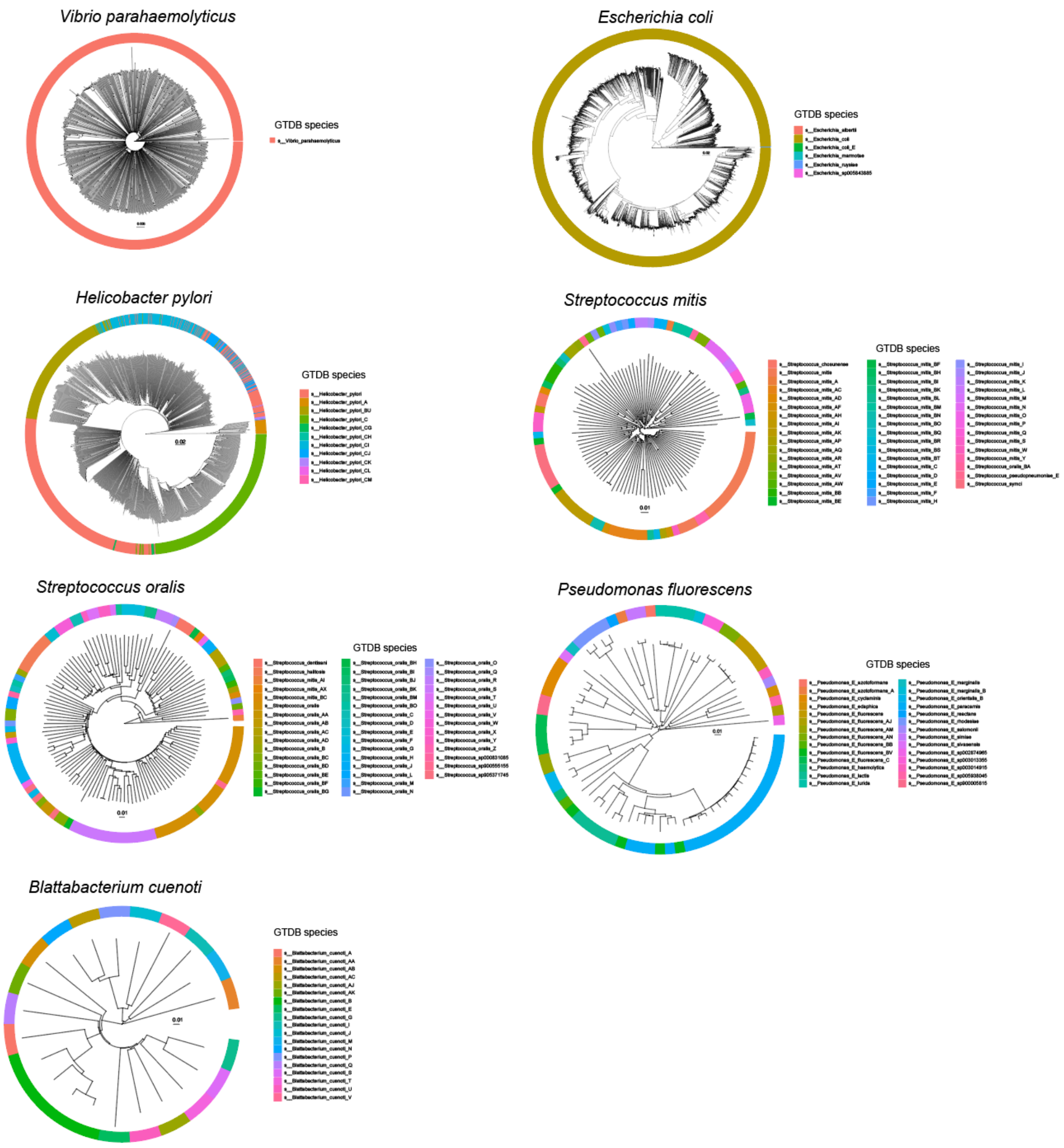

**Supplementary Figure 7.** Correlation between parameter relationships derived from NCBI and GTDB taxonomies. *R*: correlation coefficient.

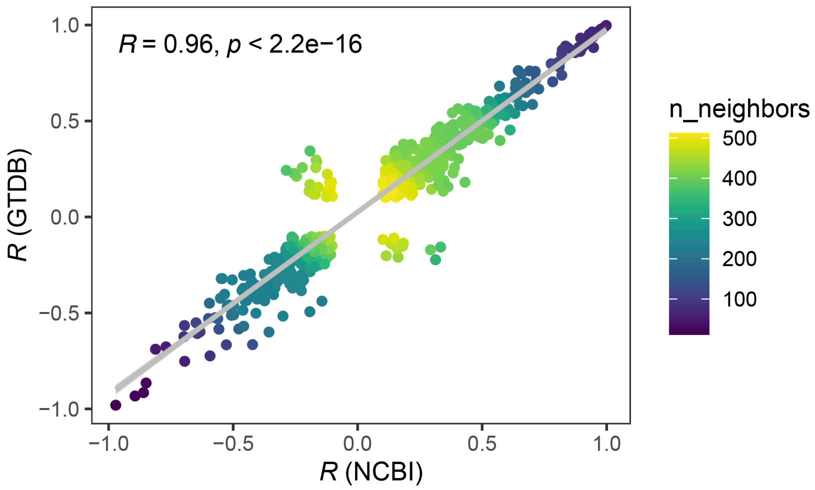

### Supplementary Text

**Linux command-line scripts for calculating genomic and population genomic parameters**

##### Genome size and GC content

```
quast.py -o <outputdir> <fasta>
```

<outputdir>: directory to store result files
<fasta>: genome sequence in FASTA format

##### Gene annotation

```
prokka --outdir <outputdir> --locustag <accession> --prefix <accession> --gcode <gcode> <fasta>
```

<outputdir>: directory to store all result files
<accession>: genome accession
<gcode>: genetic codes for each species obtained from the NCBI taxonomy database
<fasta>: genome sequence in FASTA format

##### Codon bias parameters

```
codonw <cds> -all_indices -nomenu -silent
```

<cds>: coding sequences of a genome

##### Restriction-modification (R-M) system genes

```
blastp -db <REBASE_Gold_Standards_Protein.faa> -query <faa> -outfmt "6 qacc sacc qlen length pident evalue bitscore qstart qend sstart send" -out <blastp.out> -evalue 0.01
```

<REBASE_Gold_Standards_Protein.faa>: standard protein sequences of RM system genes retrieved from REBASE
<faa>: protein sequences of each genome from Prokka output
<blastp.out>: blastp output

##### Repetitive regions

```
trf <fasta> 2 7 7 80 10 50 2000 -d -h
makeblastdb -in <fasta> -dbtype nucl
blastn -query <fasta> -db <fasta> -out <blastn.out> -outfmt 6
```

<fasta>: genome sequence in FASTA format
<blastn.out>: blastn output

##### CRISPR arrays

```
python CRISPRidentify.py --file <fasta>
```

<fasta>: genome sequence in FASTA format

**Phage sequences**

```
python PhiSpy.py <gbk> -o <outputdir>
```

<gbk>: genome sequence in GenBank format

<outputdir>: directory to store result files

**Insertion sequence**

```
isescan.py --seqfile <fasta> --output <outputdir>
```

<fasta>: genome sequence in FASTA format

<outputdir>: directory to store result files

**$\pi_s$ and *dN/dS***

```
yn00 <cds.crl>
```

<cds.ctl>: input file for PAML analysis, formatted as follows:

```
seqfile = cds.fa              * CDS alignments
outfile = dNdS.out            * main result file
verbose = 0                   * 1: detailed output; 0: concise output

icode = 0                     * 0: universal code; 1: mammalian mt; 2–10: see below

weighting = 0                 * weighting pathways between codons (0/1)
commonf3x4 = 0                * use one set of codon frequencies for all pairs (0/1)
ndata = 1
* Genetic codes: 0:universal, 1:mammalian mt., 2:yeast mt., 3:mold mt.,
* 4: invertebrate mt., 5: ciliate nuclear, 6: echinoderm mt.,
* 7: euplotid mt., 8: alternative yeast nu. 9: ascidian mt.,
* 10: blepharisma nu.
* These codes correspond to transl_table 1 to 11 of GENEBANK.
```

**Linkage disequilibrium**

```
java -jar Haploview.jar -n -info <snp.info> -pedfile <snp.ped> -maxdistance 10 -dprime -minGeno 0.6 -minMAF 0 -hwcutoff 0 -log <ld.log>
```

<snp.info>: SNP allele information required for Haploview

<snp.ped>: SNP matrix in PED format

<ld.log>: Haploview logfile

***h/m***

```
raxmlHPC-PTHREADS-SSE3 -T 10 -f x -p 1234567 -s <snp.fa> -m GTRGAMMA -n dist > <raxml.dist>
python2 calcHM_ConSpeciFix.py <raxml.dist> <snp.fa> > <hm.out>
```

<snp.fa>: concatenated SNP sequences in FASTA format
<raxml.dist>: sample distances estimated by RAxML based on SNP distances
<hm.out>: output file of the ratio of homoplasic to non-homoplasic alleles (h/m) based on the ConSpeciFix script

**Core- and pan-genome**

```
panaroo -i <*.gff> -o <outputdir> --clean-mode strict -t 10
```

<*.gff>: GFF files generated by Prokka for all representative genomes of a species
<outputdir>: directory to store result files

**Recombination-related parameter estimation based on all representative genomes**

```
mcorr-xmfa <aln.xfma> <process_prefix>
mcorrFitCompare <process_prefix> <mcorr_output_prefix>
```

<aln.xfma>: whole-genome alignment of representative genomes of a species in XMFA format
<output_prefix>: prefix of procrsss files
<mcorr_output_prefix>: prefix of mcorr-estimated recombination-related parameter values output

**Recombination-related parameter estimation of clonal group (CG)**

```
FastTree -nt <snp.fa> > <tree>
ClonalFrameML <tree> <aln> <output_prefix> -emsim 100 -ignore_user_sites <noncore.sites> > <cfml.log>
```

<snp.fa>: concatenated SNP sequences in FASTA format
<output_prefix>: prefix of output files

**Substitution rate estimation of clonal group (CG)**

```
FastTree -nt <non_rec_snp.fa> > <non_rec_tree>
Rscript BactDating.R <non_rec_tree> <snp_num> <year_info> <BactDate.tree> <BactDate.pdf> > <BactDate.log>
```

<snp.fa>: concatenated non-recombined (flited snps located in recombination regions identified by clonalframeML ) SNP sequences in FASTA format
<non_rec_tree>: phylogenetic tree construsted based on non-recombined SNPs
<snp_num>: number of non-recombined SNPs
<year_info>: isolation year information of strains
<BactDate.tree>: dated phylogenetic tree inffered by BactDating
<BactDate.pdf>: BactDating output figures
<BactDate.log>: BactDating log file

BactDating.R script as follows:

```
args <- commandArgs(trailingOnly = TRUE);

tree=args[1];
snp_num=args[2];
year=args[3];
outtree=args[4];
outfile=args[5];
outmcmc=args[6];

library(BactDating);
library(ape);
library(coda);
pdf(outfile,width=7, height=5);

t=read.tree(tree);
snp_num=as.numeric(snp_num);
t$edge.length=t$edge.length*snp_num;
d=read.table(year,header=T);
year=d$year;
names(year)=d$id;

rooted=initRoot(t,year);
root2tip=roottotip(rooted,year);

res=bactdate(t,year,nbIts=1e7,updateRoot=T,showProgress=T);
write.tree(res$tree,outtree)
plot(res,'treeCI',show.tip.label = F);

mcmc=as.mcmc.resBactDating(res);
write.table(res$CI,outmcmc);
effectiveSize(mcmc);

dev.off();
```
